# Mucoromycotina fine root endophyte fungi form nutritional mutualisms with vascular plants

**DOI:** 10.1101/531103

**Authors:** Grace A. Hoysted, Alison S. Jacob, Jill Kowal, Philipp Giesemann, Martin I. Bidartondo, Jeffrey G. Duckett, Gerhard Gebauer, William R. Rimington, Sebastian Schornack, Silvia Pressel, Katie J. Field

## Abstract

Fungi and plants have engaged in intimate symbioses that are globally widespread and have driven terrestrial biogeochemical processes since plant terrestrialisation >500 Mya. Recently, hitherto unknown nutritional mutualisms involving ancient lineages of fungi and non-vascular plants have been discovered. However, their extent and functional significance in vascular plants remains uncertain. Here, we provide first evidence of abundant carbon-for-nitrogen exchange between an early-diverging vascular plant (Lycopodiaceae) and Mucoromycotina (Endogonales) fine root endophyte regardless of changes in atmospheric CO_2_ concentration. Furthermore, we provide evidence that the same fungi also colonize neighbouring non-vascular and flowering plants. These findings fundamentally change our understanding of the evolution, physiology, interrelationships and ecology of underground plant-fungal symbioses in terrestrial ecosystems by revealing an unprecedented nutritional role of Mucoromycotina fungal symbionts in vascular plants.

## Introduction

Plant terrestrialisation >500 Mya [1] was facilitated by the formation of mutualistic symbioses with fungi, through which the earliest plants gained access to mineral nutrients in exchange for photosynthetically-fixed carbon (C) under ancient, high atmospheric CO_2_ concentrations (a[CO_2_]) [2]. It was long hypothesised that this ancient mycorrhizal-like symbiosis was closely related to and subsequently evolved into widespread modern-day arbuscular mycorrhizas (AM) formed with plant roots by Glomeromycotina fungi [3, 4]. However, recent molecular, cytological, physiological and paleobotanical evidence has strongly indicated that early fungal associates were likely to be more diverse than has previously been assumed [5–7]. Members of the earliest diverging clade of an ancient land plant lineage, Haplomitriopsida liverworts, are now known to form a[CO_2_]-responsive mycorrhizal-like associations with Mucoromycotina fungi [5, 8] which also colonise other early diverging plant lineages, namely hornworts, lycophytes and ferns [9, 10]. Mucoromycotina represent an ancient fungal lineage considered to branch earlier than, or sister to, the Glomeromycotina [11, 12], thus its recent identification in a range of modern non-vascular plants [6] and plant fossils [7, 13] supports the idea that the colonisation of Earth’s land masses by plants was facilitated not only by Glomeromycotina but also by Mucoromycotina fungal symbionts [14]. Latest discoveries of putative Mucoromycotina fungi in vascular land plants [10, 15, 16] indicate that root symbiotic versatility and diversity [17] has been grossly underestimated across extant plants.

Although Mucoromycotina fungal symbioses in non-vascular plants have, to date, received most attention [5, 6, 9, 18], there are now several reports of their occurrence in vascular plants [10, 15–17, 19]. It has been suggested that the globally widespread, arbuscule-forming fine root endophytes (FRE), classified as *Glomus tenue (*or *Planticonsortium tenue* [20]), and which occur across a wide range of vascular groups [19] are closely related to the Mucoromycotina symbionts of non-vascular plants. These findings have major ramifications for our understanding of the past and present diversity and function of plant-fungal nutritional symbioses [21], suggesting Mucoromycotina fungal symbiosis is not limited to ancient plant lineages but is in fact widespread throughout extant land plants. However, it remains unclear whether the putative Mucoromycotina FRE fungi detected in vascular plants to date are the same in terms of function and identity as the mutualistic Mucoromycotina fungal symbionts detected in non-vascular plants.

As lycophytes are considered to be the earliest divergent extant vascular plant lineage [22], the discovery of non-Glomeromycotina fungal associates in lycophyte roots and gametophytes is particularly significant. For over 100 years, the fungal associations in lycophytes have been thought of as being AM-like but with unique “lycopodioid” features [23]. However, global analysis of fungal associates in 20 lycophytes [15] has now shown their colonisation is broadly similar to that of hornworts [9], with many species forming single and/or dual associations with both Glomeromycotina arbuscular mycorrhiza fungi (AMF) and Mucoromycotina FRE fungi [15]. Remarkably, every sample of *Lycopodiella inundata* - a species common in wet habitats across the Northern Hemisphere - examined so far appears colonised exclusively by Mucoromycotina FRE fungi [15]. Since a major obstacle to studying Mucoromycotina FRE function has been finding plants that are not co-colonized by coarse root endophytes (i.e. Glomeromycotina AMF) [19], *L. inundata* provides a unique and important opportunity to dissect the symbiotic function of FRE in a vascular plant. This is particularly pertinent given that the functional significance of Mucoromycotina FRE associations in vascular plants and their response to changing a[CO_2_] relevant to conditions during the Paleozoic era and the time of vascular plant divergence are completely unknown [17, 19]. Indeed, there is no evidence of nutritional mutualism between any vascular plant and Mucoromycotina FRE [17].

Here, we address these critical knowledge gaps by investigating the cytology, function and identity of the fungal association in *L. inundata (*Figure 1 a, b) under simulated ancient and modern a[CO_2_]. We use a combination of molecular biology, radio- and stable isotope tracers, and cytological analyses to address the following questions:

1. Do Mucoromycotina fungal symbionts of *L. inundata* co-occur in neighbouring angiosperm roots and non-vascular plant rhizoids?
2. Are there characteristic cytological signatures or features of Mucoromycotina fungal associations in *L. inundata* compared to those formed in non-vascular plants?
3. What is the function of Mucoromycotina fungal associations in lycophytes in terms of carbon-for-nutrient exchange and is it affected by a[CO_2_]?

**Figure 1.**
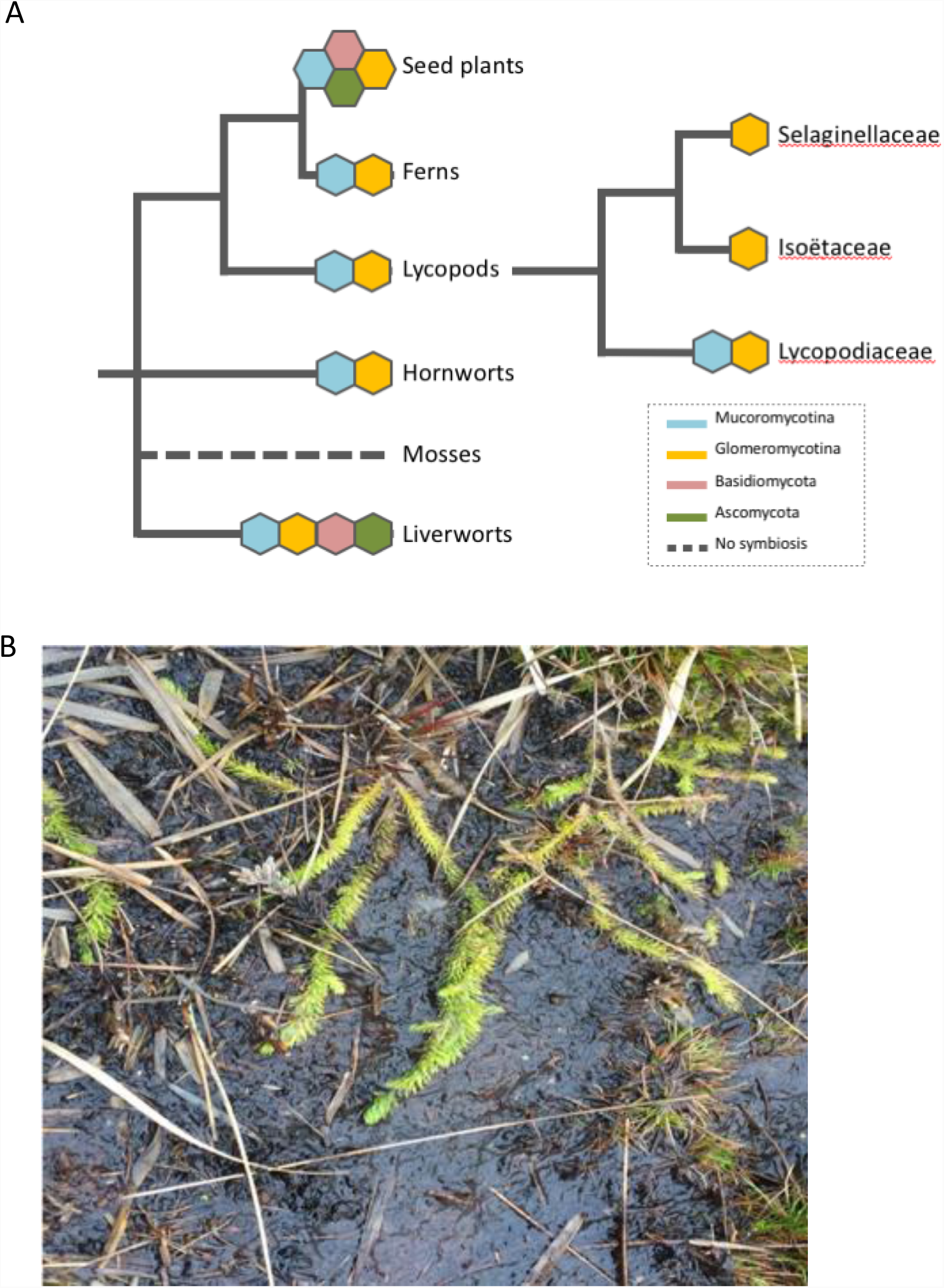
Land plant phylogeny and species used in the present study. (a) Land plant phylogeny showing key nodes alongside commonly associated fungal symbionts [6, 11, 23, 31] (b) *Lycopodiella inundata* at Thursley Common, Surrey, June 2017.

## Methods

### Plant material

*Lycopodiella inundata (*L.), neighbouring angiosperms (the grasses *Holcus lanatus, Molinia caerulea* and the rush *Juncus bulbosus*), and liverworts (*Fossombronia foveolata*) were collected from Thursley National Nature Reserve, Surrey, UK (SU 90081 39754) in June 2017. The *L. inundata* plants were planted directly into pots (90 mm diameter x 85 mm depth) containing acid-washed silica sand. Soil surrounding plant roots was left intact and pots were weeded regularly to remove other plant species. The other plants collected in Thursley, and additional plants from three other UK field sites (Supplementary Table S1), were used for cytological and molecular analyses. Additional vascular plants from Thursley were used for stable isotope analyses.

### Growth conditions

Based on the methods of Field *et al.* [8], three windowed cylindrical plastic cores covered in 10 µm nylon mesh (Supplementary Figure S1) were inserted into the substrate within each experimental pot. Two of the cores were filled with a homogenous mixture of acid-washed silica sand, compost (Petersfield No.2, Leicester, UK) and native soil gathered from around the roots of wild plants (in equal parts making up 99% of the core volume) and fine-ground tertiary basalt (1% core volume) [8]. The third core was filled with glass wool to allow below-ground gas sampling throughout the ^14^C-labelling period to monitor soil community respiration.

The *L. inundata* plants were maintained in controlled environment chambers (Micro Clima 1200, Snijders Labs, The Netherlands). Plants were grown at two contrasting CO_2_ atmospheres; 440 ppm a[CO_2_] to represent a modern-day atmosphere, or at 800 ppm a[CO_2_] to simulate Paleozoic atmospheric conditions on Earth at the time vascular plants are thought to have diverged [2]. a[CO_2_] was monitored using a Vaisala sensor system (Vaisala, Birmingham, UK), maintained through addition of gaseous CO_2_. Cabinet settings and contents were alternated every four weeks, and all pots were rotated within cabinets to control for cabinet and block effects. Plants were acclimated to chamber/growth regimes (see Supplementary Information) for four weeks to allow establishment of mycelial networks within pots and confirmed by hyphal extraction from soil and staining with trypan blue [24]. Additionally, roots were stained with acidified ink for the presence of fungi, based on the methods of Brundrett *et al.* [24].

### Molecular identification of fungal symbionts

All plants (Supplementary Table S1) were processed for molecular analyses within one week of collection. Genomic DNA extraction and purification from all specimens and subsequent amplification, cloning and sequencing were performed according to methods from Rimington *et al*. [10]. The fungal 18S ribosomal rRNA gene was targeted using the broad specificity fungal primer set NS1/EF3 and a semi-nested approach with Mucoromycotina- and Glomeromycotina-specific primers described in Desirò *et al*. [9] for the experimental *L. inundata* plants and all other field collected plant material using Mucoromycotina-specific primers. Resulting partial 18S rDNA sequences were edited and preliminarily identified with BLAST in Geneious v. 8.1.7 [25]. Chimeric sequences were detected using the UCHIME2 algorithm [26] in conjunction with the most recent non-redundant SSU SILVA database (SSU Ref NR 132, December 2017, www.arb-silva.de). Sequences identified as Mucoromycotina sp. were aligned with MAFFT prior to removing unreliable columns using the default settings in GUIDANCE2 (http://guidance.tau.ac.il). The best-fit nucleotide model for phylogenetic analysis was calculated using Smart Model Selection [27]. Maximum Likelihood (ML) with 1,000 replicates was performed using PhyML 3.0 [28]. Bayesian inference analysis was conducted in Mr Bayes version 3.2.6 [29] with four Markov chain Monte Carlo (MCMC) strands and 10^6^ generations. Consensus trees were produced after excluding an initial burn-in of 25% of the samples (Supplementary Figures S2-8). Representative DNA sequences were deposited in GenBank.

### Cytological analyses

*Lycopodiella inundata* gametophytes, young sporophytes (protocorms) and roots of mature plants (both wild and experimental), roots of angiosperms (*Holcus lanatus, Molinia caerulea* and *Juncus bulbosus*), and liverwort gametophytes (*Fossombronia foveolata*) were either stained with trypan blue [24], which is common standard for identifying FRE [19], and photographed under a Zeiss Axioscope (Zeiss, Oberkochen, Germany) equipped with a MRc digital camera, or processed for scanning electron microscopy (SEM) within 48 hrs of collection [30]. For SEM we followed the protocol by Duckett *et al.* [31] (see Supplementary Information). For experimental plants of *L. inundata*, ten randomly selected roots per treatment were cut into up to six segments (depending on root length) and colonization by FRE scored as absent or present for each segment under the SEM.

### Quantification of C, ^33^P and ^15^N fluxes between lycophytes and fungi

After the four-week acclimation period, 100 µl of an aqueous mixture of ^33^P-labelled orthophosphate (specific activity 111 TBq mmol^−1^, 0.3 ng ^33^P added; Hartmann analytics, Braunschweig, Germany) and ^15^N-ammonium chloride (1mg ml^−1^; 0.1 mg ^15^N added; Sigma, Dorset, UK) was introduced into one of the soil-filled mesh cores in each pot through the installed capillary tube (Supplementary Figure S9a). In half (12) of the pots, cores containing isotope tracers were left static to preserve direct hyphal connections with the lycophytes. Fungal access to isotope tracers was limited in the remaining half (12) of the pots by rotating isotope tracer-containing cores through 90°, thereby severing the hyphal connections between the plants and core soil. These were rotated every second day thereafter, thus providing a control treatment that allows us to distinguish between fungal and microbial contributions to tracer uptake by plants. Assimilation of ^33^P tracer into above-ground plant material was monitored using a hand-held Geiger counter held over the plant material daily.

At detection of peak activity in above-ground plant tissues (21 days after the addition of the ^33^P and ^15^N tracers), the tops of ^33^P and ^15^N-labelled cores were sealed with plastic caps and anhydrous lanolin and the glass wool cores were sealed with rubber septa (SubaSeal, Sigma, Dorset, UK). Each pot was sealed into a 3.5 L, gas-tight labelling chamber and 2 ml 10% lactic acid was added to 30 µl NaH^14^CO_3_ (specific activity 1.621 GBq/mmol^−1^; Hartmann Analytics, Braunschweig, Germany) in a cuvette within the chamber prior to cabinet illumination at 0800 (Supplementary Figure S9b), releasing a 1.1-MBq pulse of ^14^CO_2_ gas. Pots were maintained under growth chamber conditions, and 1 ml of gas was sampled after 1 hour and every 1.5 hours thereafter. Below-ground respiration was monitored via gas sampling from within the glass-wool filled core after 1 hour and every 1.5 hours thereafter for ∼16 h.

### Plant harvest and sample analyses

Upon detection of maximum below-ground flux of ^14^C, plant materials and soil were separated, freeze-dried, weighed and homogenised. The ^33^P activity in plant and soil samples was quantified by digesting in concentrated H_2_SO_4_ (see Supplementary Information) and liquid scintillation (Tricarb 3100TR liquid scintillation analyser, Isotech, Chesterfield, UK). The quantity of ^33^P tracer that was transferred to the plant by it’s fungal partner was then calculated using previously published equations [32] (see Supplementary Information). Total ^33^P in plants without access to the tracer through core rotation (i.e. assimilated through alternative soil microbial P-cycling processes and/or diffusion from core) was subtracted from the total ^33^P in plants with access to the core contents via intact fungal hyphal connections to give fungal acquired ^33^P.

Between two and four mg of freeze-dried, homogenised plant tissue was weighed into 6 x 4 mm^2^ tin capsules (Sercon Ltd. Irvine, UK) and ^15^N abundance was determined using a continuous flow IRMS (PDZ 2020 IRMS, Sercon Ltd. Irvine, UK). Air was used as the reference standard, and the IRMS detector was regularly calibrated to commercially available reference gases. The ^15^N transferred from fungus to plant was then calculated using equations published previously [18] (see Supplementary Information). Total ^15^N in plants without access to the isotope because of broken hyphal connections between plant and core contents was subtracted from total ^15^N in plants with intact hyphal connections to the mesh-covered core to give fungal-acquired ^15^N.

The ^4^C activity of plant and soil samples was quantified through sample oxidation (307 Packard Sample Oxidiser, Isotech, Chesterfield, UK) followed by liquid scintillation. Total C (^12^C + ^14^C) fixed by the plant and transferred to the fungal network was calculated as a function of the total volume and CO_2_ content of the labelling chamber and the proportion of the supplied ^14^CO_2_ label fixed by plants (see Supplementary Information). The difference in total C between the values obtained for static and rotated core contents is considered equivalent to the total C transferred from plant to symbiotic fungus within the soil core, noting that a small proportion will be lost through soil microbial respiration. The total C budget for each experimental pot was calculated using equations from Cameron *et al*. [33] (see Supplementary Information).

### Stable isotope signatures of neighbouring plants

*Lycopodiella inundata* and *J. bulbosus* were collected from Thursley National Nature Reserve, Surrey, together with co-occurring reference plants from six 1 m^2^ plots in May 2018, following the sampling scheme of Gebauer and Meyer [34]. Five plant species representing three different types of mycorrhizal associations served as reference plants: two ericoid mycorrhizal species (*Erica tetralix*, collected on six plots; *Calluna vulgaris*, collected on three plots), two ectomycorrhizal species (*Pinus sylvestris* and *Betula pendula* seedlings, both from one plot) and one arbuscular mycorrhizal species (*Molinia caerulea* from six plots). Relative C and N isotope natural abundances of dried and ground leaf and root samples were measured in a dual element analysis mode with an elemental analyser (Carlo Erba Instruments 1108, Milan, Italy) coupled to a continuous flow isotope ratio mass spectrometer (delta S, Finnigan MAT, Bremen, Germany) via a ConFlo III open-split interface (Thermo Fisher Scientific, Bremen, Germany) as described in Bidartondo *et al*. [35]. Relative isotope abundances (δ values) were calculated, calibrated and checked for accuracy using methods detailed in Supplementary Information.

### Statistics

Effects of plant species, a[CO_2_] and the interaction between these factors on the C, ^33^P and ^15^N fluxes between plants and Mucoromycotina fungi were tested using analysis of variance (ANOVA) or Mann-Whitney U where indicated. Data were checked for homogeneity and normality. Where assumptions for ANOVA were not met, data were transformed using log_10_. If assumptions for ANOVA were still not met, a Mann Whitney U statistical test was performed. All statistics were carried out using the statistical software package SPSS Version 24 (IBM Analytics, New York, USA). Stable isotope patterns were tested for normality and equal variance. If the requirements of parametric data and equal variance were fulfilled, one-way ANOVA was applied, while for non-parametric data Kruskal-Wallis tests were performed. Leaves and roots were tested separately. Mean values are given with standard deviations.

## Results

### Molecular identification of fungal symbionts

Analysis of experimental *L. inundata* plants grown under ambient and elevated a[CO_2_] confirmed that they were colonised by Mucoromycotina fungi. Glomeromycotina sequences were not detected. Mucoromycotina OTUs were detected before and after the experiments (Supplementary Figure S2); these same OTUs had previously been identified in wild-collected lycophytes from diverse locations [10].

Diverse and shared Mucoromycotina fungi OTUs were detected in wild *L. inundata*, liverworts and angiosperms growing adjacently in the same UK locations (Supplementary Table S2, Fig. S2-8) in the following combinations: *L. inundata, F. foveolata, M. caerulea* and *J. bulbosus (*Thursley Common, Surrey); *L. inundata, F. foveolata* and *J. bulbosus (*Norfolk); *F. foveolata* and *H. lanatus (*Lynn Crafnant, Wales). Mucoromycotina OTUs were also detected in *L. inundata* from Studland Heath, Dorset.

### Cytology of fungal colonisation in plants

Trypan blue staining and SEM revealed two distinct fungal symbiont morphologies consisting of either coarse hyphae (>3 µm diameter) and large vesicles (>20 µm diameter) or fine branching hyphae (<2 µm diameter) with small swellings/vesicles (usually 5-10 but up to 15 µm diameter) (Figures 2-3). Both morphologies were observed in the gametophyte of the liverwort *F. foveolata (*Figures 2a, b, 3a; Supplementary Figure S10), in the roots of the grasses *H. lanatus (*Figure 2f) and *M. caerulea (*Figure 2g, h), and the rush *J. bulbosus (*Figure 3h, i). In the roots of wild and experimental plants of *L. inundata*, only fine hyphae were detected (Figures 2c-e, 3f, g). As in the other plants analysed, these fine hyphae were aseptate and formed both intercalary and terminal swellings/vesicles but, in contrast to the grasses, never arbuscules (Supplementary Figure S10). Similar fungal morphology was also observed in protocorm cells of newly developing sporophytes (Figure 3b, c) and in gametophytes of *L. inundata (*Supplementary Figure S11). However, in these early developmental stages, fungal colonization exhibits a distinct zonation: an outer intracellular zone and a more central, strictly intercellular zone (Figure 3d, e; Supplementary Figure S11b, c, g). In the intracellular zone, fungal colonization is the same as in the sporophyte roots and consists of fine hyphae with intercalary and terminal swellings/vesicles (Figure 3b, c; Supplementary Figure S11i). Unique to the gametophyte generation, in the outermost cortical layers, the fungus also forms tightly woundcoils (hyphae up to 2.5 µm in diameter) with larger vesicles (15-20 µm) (Supplementary Figure S11d), as described before in *Lycopodium clavatum* [36]. Both gametophyte and early developmental stages of the sporophyte generation develop a conspicuous central system of large, mucilage-filled intercellular spaces. In this region, the fungus becomes strictly intercellular (Figure 3d, e; Supplementary, Figure S11g). The intercellular hyphae are initially fine and with small swellings/vesicles (Figure 3d, Supplementary Figure S11e), as their intracellular counterparts, but soon enlarge and eventually reach diameters in excess of 3 µm (Supplementary Figure S11f), with no swellings/vesicles present at this stage. While no morphological differences were detected between fungal root associates of the two experimental *Lycopodiella* grown under contrasting a[CO_2_], those grown under 800 ppm a[CO_2_] had more colonization (44 out of 56 root segments; 79%) than those grown under 440 ppm a[CO_2_] (31 out of 58 root segments; 53%).

**Figure 2.**
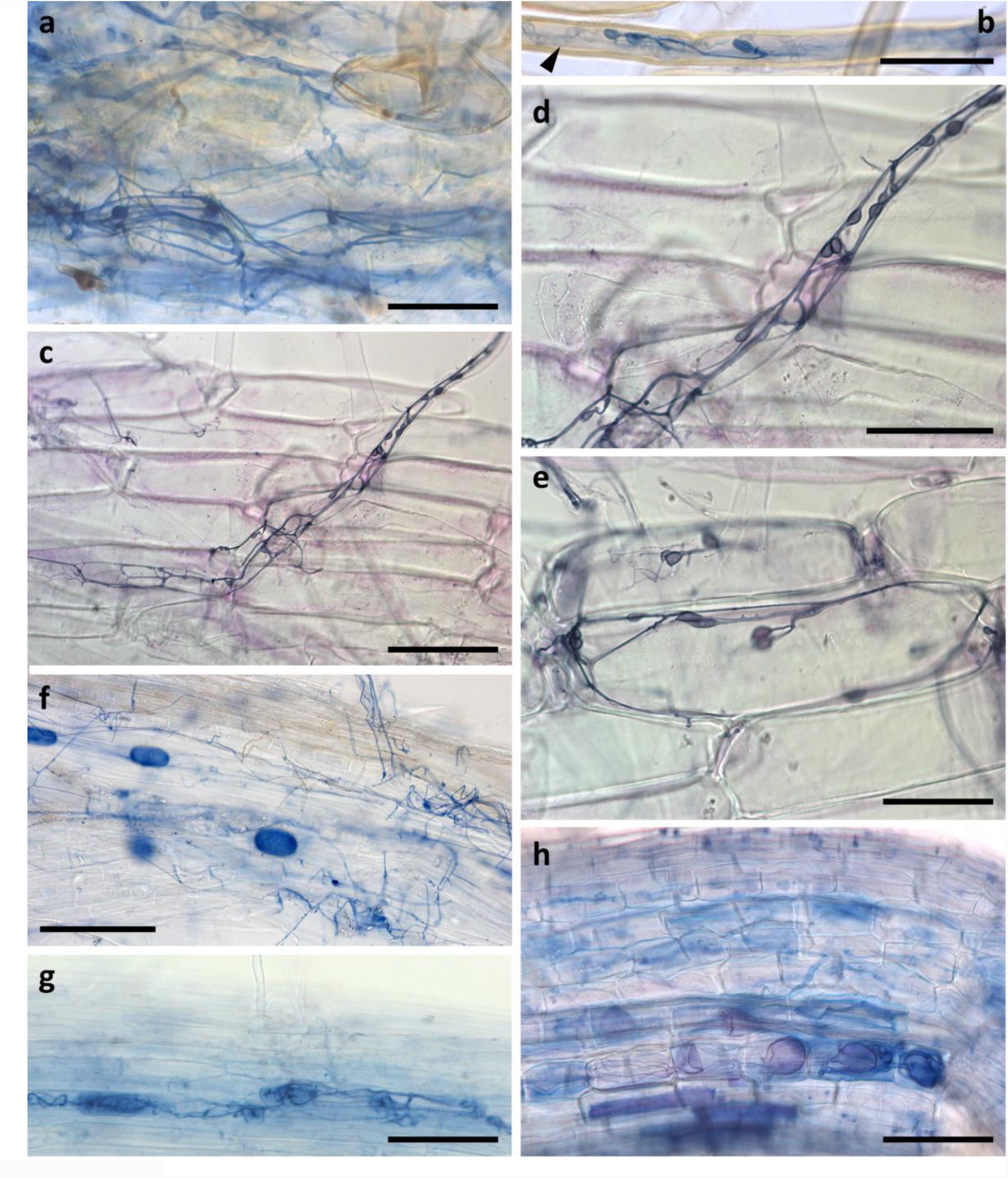
Light micrographs of trypan blue stained tissues. (a) Branching fine hyphae with small swellings/vesicles in thallus cells and rhizoid (b) of the liverwort *Fossombronia foveolata* colonized by both Mucoromycotina fine root endophytes (FRE) and Glomeromycotina, in (b) also note the coarse hyphae (arrowhead). (c-e) Fine hyphae with small swellings/vesicles in the root hairs and root cells of the lycophyte *Lycopodiella inundata* colonized by Mucoromycotina FRE only. (f) Fine hyphae with small swellings/vesicles and large vesicles in a root of the grass *Holcus lanatus* colonized by both Mucoromycotina FRE and Glomeromycotina. (g-h) Roots of the grass *Molinia caerulea* colonized by both Mucoromycotina FRE and Glomeromycotina, showing fine hyphae (g) and coarse hyphae with large vesicles (h). Scale bars: (a, b, d-f) 50 µm, (c, g, h) 100 µm.

**Figure 3.**
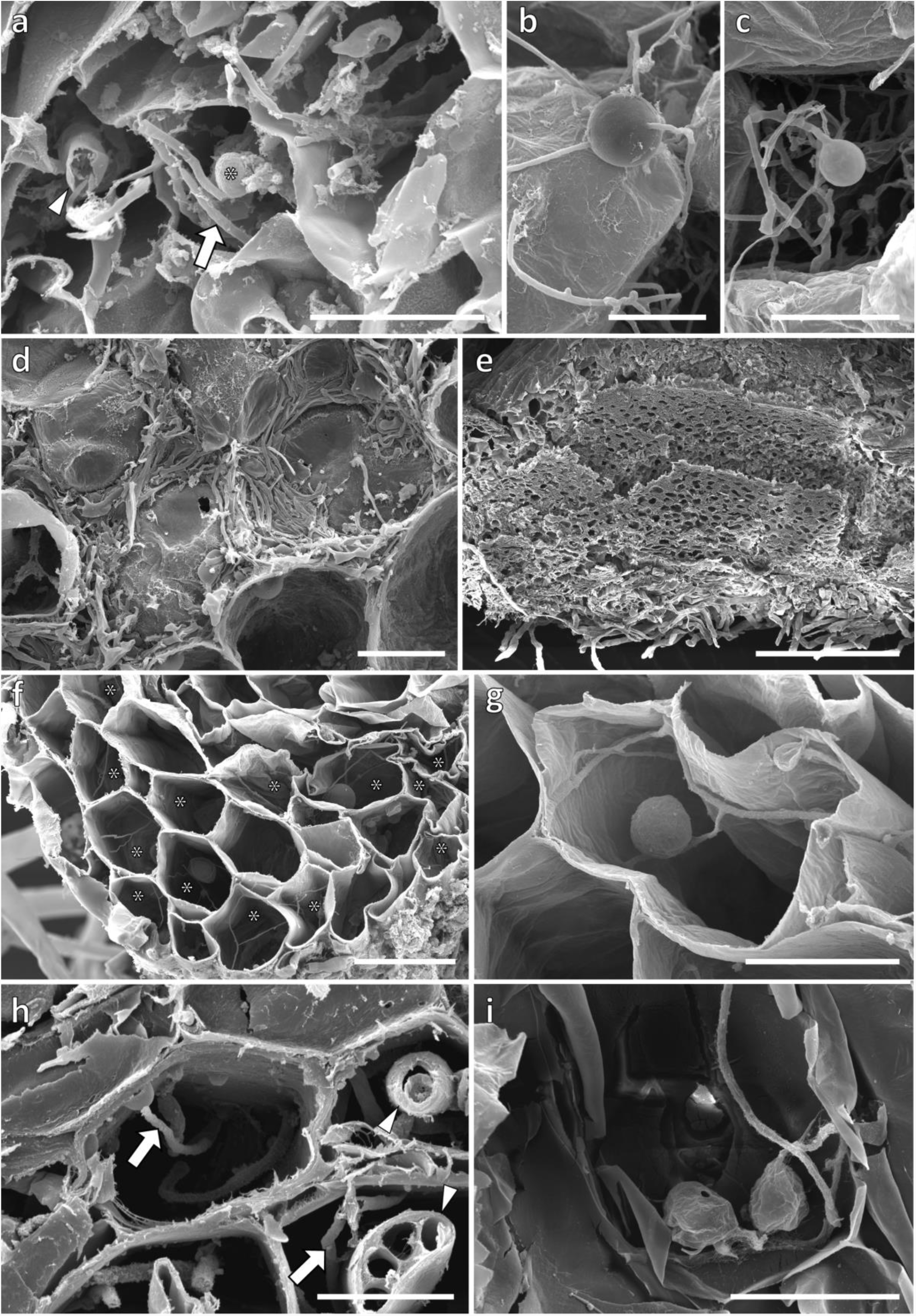
Scanning electron micrographs. (a) Fine hyphae (arrows) with a small swelling/vesicle (*) in the thallus cells of *Fossombronia foveolata*, also note the much coarser hyphae (arrowheads). (b-g) Fungal colonization in *Lycopodiella inundata*. (b, c) Intercalary (b) and terminal (c) small swellings/vesicles on fine hyphae in the ventral cell layers of a protocorm. Centrally and above this intracellular colonization zone, the fungus becomes exclusively intercellular (d, e). (f, g) Cross sections of roots showing branching fine hyphae with small swellings/vesicles. (h, i) Cross sections of roots of *Juncus bulbusus* showing fine (arrow) and coarse (arrowheads) hyphae (h) and a fine hypha with small swellings/vesicles (i). Scale bars: (e) 500 µm, (f) 50 µm, (a, d, g, i) 20 µm, (b, c, h) 10 µm.

### Lycophyte-to-fungus C transfer

Unlike in non-vascular plants, carbon allocation to fungal symbionts by *L. inundata* were not significantly affected by a[CO_2_]. However, there were trends in line with previous findings in liverworts; *L. inundata* allocated ca. 2.8 times more photosynthate to Mucoromycotina fungi under the simulated Paleozoic a[CO_2_] of 800 ppm (Figure 4a) compared with plants that were grown under ambient a[CO_2_] of 440 ppm (Figure 4a; Mann-Whitney U = 194, *P =* 0.864, n = 20). In terms of total C transferred from plants to Mucoromycotina, a similar trend was observed (Figure 4b) with *L. inundata* transferring ca. 2.7 times more C to Mucoromycotina fungal partners at elevated a[CO_2_] concentrations of 800 ppm compared to those maintained under a[CO_2_] of 440 ppm (Figure 4b; Mann-Whitney U = 197.5, *P =* 0.942, n= 20).

**Figure 4.**
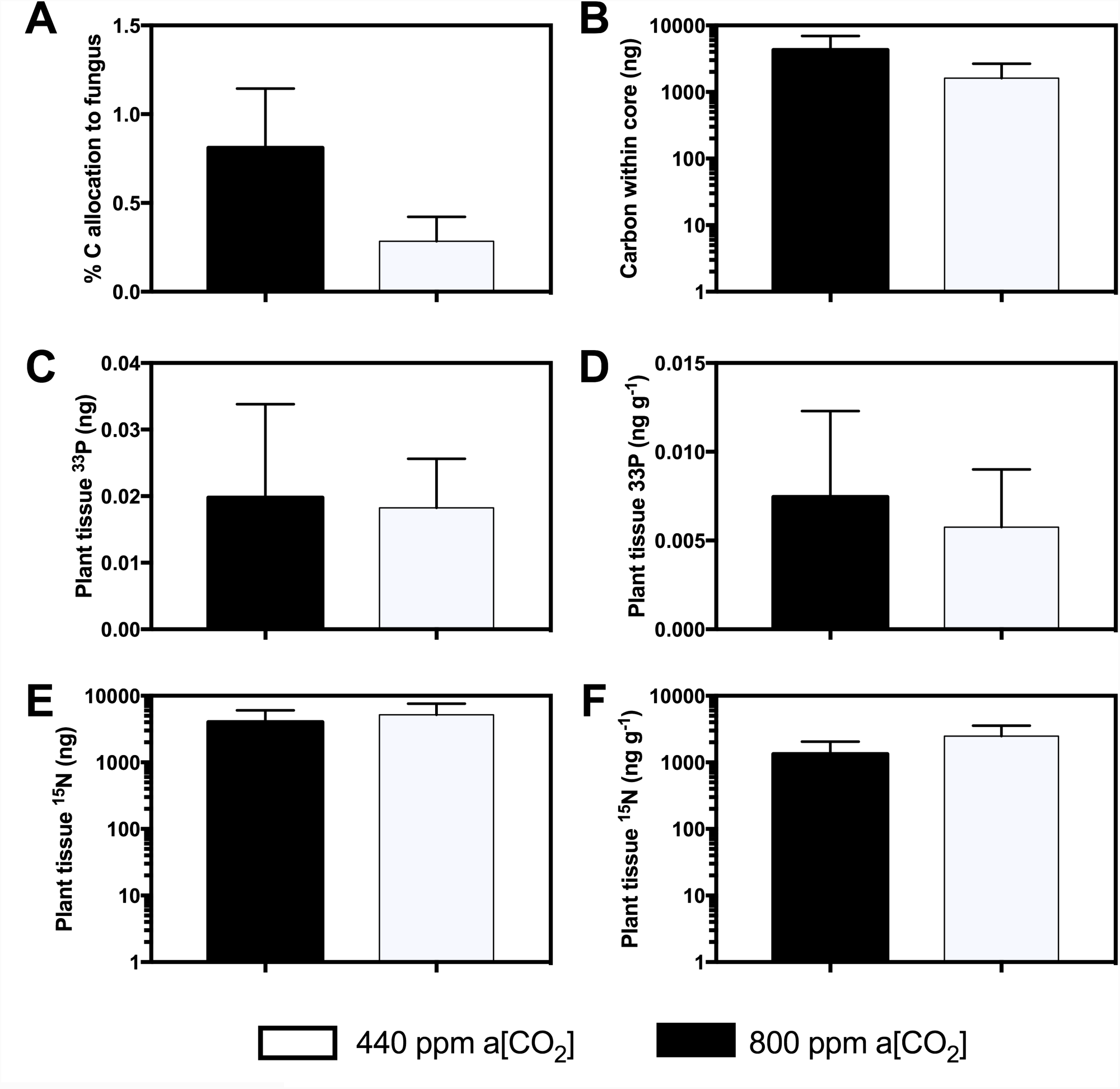
Carbon-for-nutrient exchange between *Lycopodiella inundata* and Mucoromycotina fine root endophyte fungi. **(a)** % allocation of plant-fixed C to Mucoromycotina fungi (**b)** Total plant-fixed C transferred to Mucoromycotina fungi by *L. inundata*. (**c)** Total plant tissue ^33^P content (ng) and (**d)** tissue concentration (ng g^−1^) of fungal-acquired ^33^P in *L. inundata* tissue (**e)** Total tissue ^15^N content (ng) and (**f)** concentration (ng g^−1^) of fungal-acquired ^15^N in *L. inundata* with exclusive Mucoromycotina fungal associations. All experiments conducted at a[CO_2_] of 800 p.p.m. (black bars) and 440 p.p.m. (white bars). All bars in each panel represent the difference in isotopes between the static and rotated cores inserted into each microcosm. In all panels, error bars denote SEM. In panels a and b *n* = 20 for both a[CO_2_]. In panels c-f *n* = 12 for both 800 p.p.m and 440 p.p.m a[CO_2_].

### Fungus-to-lycophyte ^33^P and ^15^N transfer

Mucoromycotina fungi transferred ^33^P and ^15^N to their plant hosts (Figure 4c-f). There were no significant differences in the amounts of either ^33^P or ^15^N tracer acquired by Mucoromycotina in *L. inundata* plant tissue when grown under elevated a[CO_2_] of 800 ppm compared to plants grown under a[CO_2_] conditions of 440 ppm, either in terms of absolute quantities (Figure 4c, ANOVA [*F*_1, 23_ = 0.009, *P =* 0.924, n = 12]; Figure 4e, ANOVA [F_1, 22_ = 0.126, *P* = 0.726, n = 12]) or when normalised to plant biomass (Figure 4d, ANOVA [F_1, 23_ = 0.085, *P* = 0.774, n = 12] and Figure 4f, ANOVA [F_1, 22_= 0.770, *P* = 0.390, n = 12]).

### Natural abundance δ^13^C and δ^15^N stable isotope signatures of plants

All leaf δ^13^C values ranged between −26.2 and −30.1 ‰ and root δ^13^C values between −24.5 and −28.9 ‰, while leaf δ^15^N values ranged from 3.3 to −10.0 ‰ and root δ^15^N values from 3.1 to −5.9 ‰ (Figure 5). Leaves of the three groups, *L. inundata (*n = 6), *J. bulbosus (*n = 6) and reference plants (n = 17), were significantly different in δ^13^C (*H*(2) = 8.758; *p* = 0.013) and δ^15^N (*H*(2) = 21.434; *P* < 0.001, Figure 5a). *L. inundata* leaves were significantly depleted in ^13^C compared to *J. bulbosus* leaves (*Q* = 2.644, *P* < 0.05) and a significant depletion of *L. inundata* leaves compared to reference plant leaves (*Q* = 2.662, *P* < 0.05, Figure 5a). The *J. bulbosus* leaves were not significantly different from reference plants in δ^13^C. No significant difference was discovered for δ^15^N in *L. inundata* and *J. bulbosus* leaves (*Q* = 1.017, *P* > 0.05), while leaves of both species were significantly enriched in ^15^N compared to the reference plants (*Q* = 2.968, *P* < 0.05; *Q* = 4.205, *P* < 0.05, Figure 5a). For the roots, only δ^15^N showed significant differences between the three groups under comparison (*F*(2) = 34.815; *P* < 0.001, Figure 5b). The *L. inundata* and *J. bulbosus* roots were not significantly distinguished in δ^15^N, however, roots of both species were significantly enriched in ^15^N compared to reference plant roots (*q* = 10.109, *p* < 0.001; *q* = 8.515, *p* < 0.001, Figure 5b).

**Figure 5.**
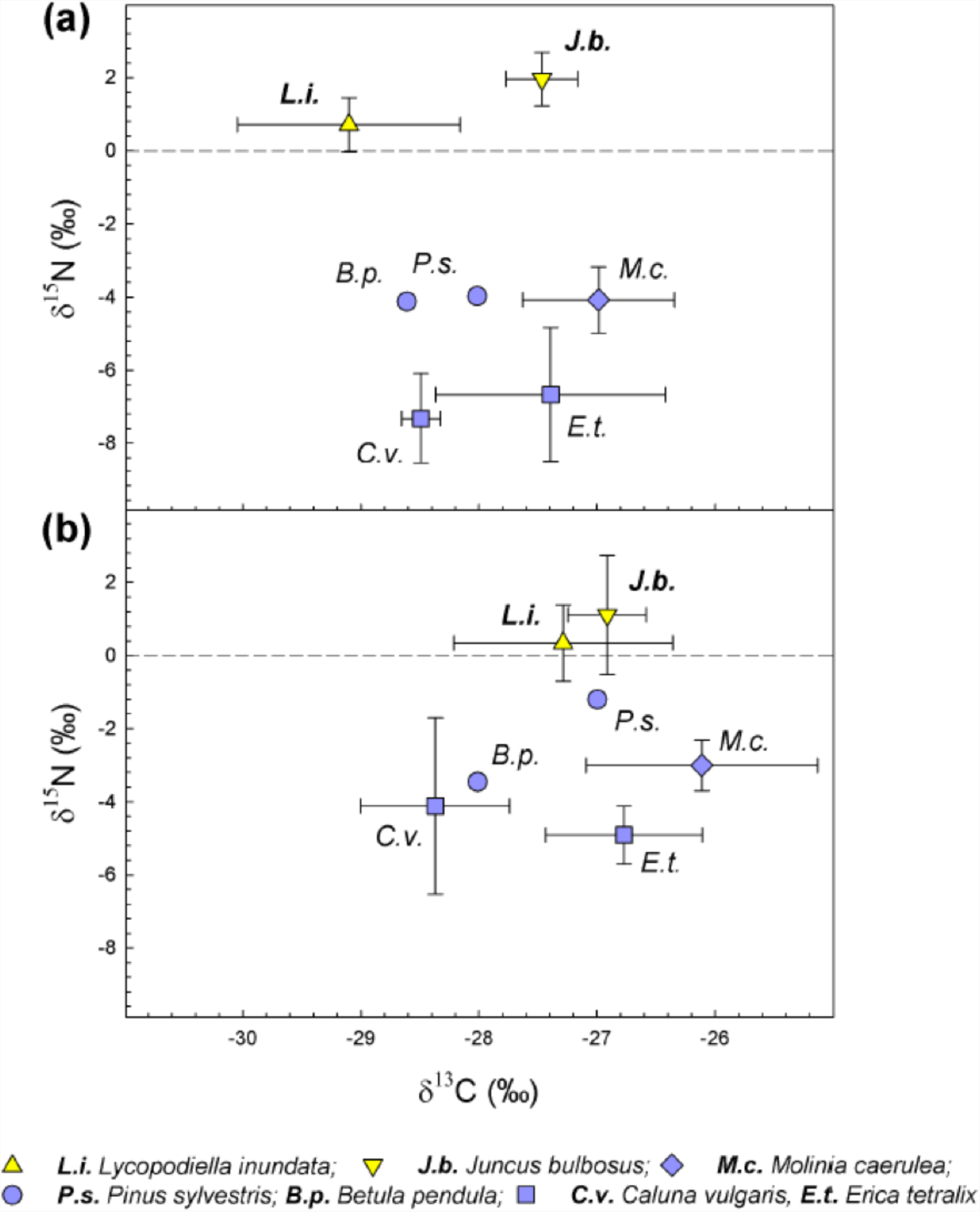
Carbon and nitrogen stable isotope natural abundance of *Lycopodiella inundata* and surrounding angiosperms. **(a)** Data from leaves. (**b)** Data from roots.

## Discussion

Our results show that the symbiosis between *L. inundata* and Mucoromycotina FRE is nutritionally mutualistic, with the fungus gaining plant-fixed C and the plant gaining fungal-acquired nutrients (Figure 4a-f). Cytological analyses of the fungus colonising the roots of *L. inundata* revealed a characteristic morphology consisting of fine, aseptate branching hyphae with terminal and intercalary swellings/vesicles. This morphology matches that described previously in a range of angiosperms colonized by FRE [16, 19] and here in grasses, a rush and a liverwort, all harbouring fungi identified molecularly as Mucoromycotina. Thus, our results provide compelling evidence for Mucoromycotina FRE being shared by plants occupying key nodes in the land plant phylogeny - from early liverworts and vascular lycophytes to the later diverging angiosperms - and that this association represents a nutritional mutualism as much in vascular as in non-vascular plants [5, 18].

Our findings raise novel and important questions regarding the evolution of mycorrhizal associations and the nature of widespread Mucoromycotina FRE fungal symbioses: what role did these fungi play during the greening of the Earth >500 Ma? How have these associations persisted and why are they so widespread today? We can now begin to address these questions with the demonstration that a vascular plant assimilates significant amounts of Mucoromycotina FRE-acquired ^15^N tracer, suggesting a significant role for Mucoromycotina FRE in vascular plant nitrogen uptake, facilitating their persistence across nearly all land plant lineages.

### Costs and benefits of hosting Mucoromycotina fungi

The amount of C transferred from *L. inundata* to Mucoromycotina symbionts was not significantly affected by a[CO_2_] (Figure 4a, b), with the fungi maintaining C assimilation across the a[CO_2_] treatments, despite colonisation being more abundant within the roots of plants grown under the elevated a[CO_2_]. Previous studies [5, 18] have demonstrated Haplomitriopsida liverwort-Mucoromycotina FRE nutritional mutualisms were affected by a[CO_2_], with the fungi gaining more C from their host liverworts under elevated a[CO_2_]. Although these experiments were carried out at higher a[CO_2_] concentrations (1,500 ppm) than the present study (800 ppm), both *Haplomitrium gibbsiae* and *Treubia lacunosa* transferred approximately 56 and 189 times less photosynthate, respectively, to their fungi [5, 18] under elevated a[CO_2_] compared to *L. inundata (*Supplementary Table S3). This trend is consistent with previous observations in vascular plants with Glomeromycotina AM [8]. When compared to other vascular plant-Glomeromycotina fungal symbioses in similar experimental systems [8], it is clear that the relative C “cost” of maintaining Mucoromycotina fungal symbionts is at least on a par with, if not greater than, that of maintaining Glomeromycotina fungi.

Lycophytes are a significant node in land plant phylogeny, widely considered as a diversification point in the mid-Paleozoic (480-360 Ma) characterised by the evolution of roots, leaves and associated vasculature [22]. The significant depletion of ^13^C observed in the leaves of *L. inundata (*Figure 5) is unlikely to be related to C gains from its Mucoromycotina fungal symbiont [37]; rather it may indicate that *L. inundata* regulate their stomata differently from *J. bulbosus* or the reference plants tested, as δ^13^C in tissues of terrestrial plants may be driven by the water use efficiency of the plant [38]. Alongside increased capacity for C capture and fixation, it is likely that increasing structural complexity in land plants across evolutionary time resulted in greater plant nutrient demand.

Glomeromycotina AM are associated with facilitation of plant P uptake and occur commonly in soils with low P availability [39, 40]. We observed no difference in the amount of fungal-acquired ^33^P tracer that was transferred to *L. inundata* sporophytes when a[CO_2_] was changed (Figure 4c, d). Given that *L. inundata* can regulate and maintain CO_2_ assimilation and thus C fixation through stomata and vasculature, it is possible that a lower a[CO_2_] would affect transfer of plant-fixed C to symbiotic fungi less than it might do in poikilohydric liverworts. The amount of ^33^P transferred to *L. inundata* plants was much less than has previously been recorded for Mucoromycotina-associated liverworts [18] or for Glomeromycotina-associated ferns and angiosperms [8], despite the same amount of ^33^P being made available, suggesting that Mucoromycotina fungi may not play a critical role in lycophyte P nutrition. Our results contrast with the view that FRE enhance plant P uptake, at least in soils with very low P [19, 41], raising questions regarding the role of FRE in *L. inundata* given that they represent a significant C outlay. Previous experiments with Mucoromycotina-associated liverworts suggest there is a role for the fungus in plant N nutrition [14, 18].

Nitrogen is an essential element for plants that is available in soils in plant-inaccessible organic forms and as plant-accessible inorganic nitrate and ammonium [42]. Our results show that although there was no significant difference in the amount of ^15^N transferred from Mucoromycotina to *L. inundata* under elevated or ambient a[CO_2_] (Figure 4e, f), up to 391 times more ^15^N was transferred to *L. inundata* than in Haplomitriopsida liverworts in comparable experiments (Supplementary Table S3) [18]. We also show that *L. inundata* and *J. bulbosus* were significantly ^15^N enriched in comparison to co-occurring reference plants with different mycorrhizal partners (Figure 5). This further supports our experimental finding that Mucoromycotina symbionts play a significant role in host plant N nutrition.

Some AM fungi transfer N to their associated hosts [43]; however, the ecological relevance of AM-facilitated N uptake is widely debated, in particular the amounts of N transferred to hosts compared to the overall N requirements of the plant [44]. Different mycorrhizal associations, i.e. ecto-, ericoid and arbuscular mycorrhizas, can influence plant δ^15^N [45]. While this distinction in N isotope abundance between plants with different mycorrhizas is almost or completely lost in conditions of higher N isotope availibility [34], it becomes significantly different under severe N limitation [46]. Exclusive plant-Mucoromycotina FRE symbioses seem to be rare, having been reported before only in the earliest-diverging Haplomitriopsida liverworts [6, 14], while all other plants, including other lycophytes [10], that form associations with these fungi appear able to do so also with Glomeromycotina, often simultaneously [14]. It is possible that the major input to *Lycopodiella* N nutrition and minor contribution to P nutrition by Mucoromycotina FRE reflect such a specialised relationship considering heathland habitats have very low plant-available N. Nevertheless, our present data combined with previous demonstrations of N transfer in liverwort-Mucoromycotina symbioses [14, 18] and emerging evidence that Mucoromycotina FRE, but not Glomeromycotina, are able to transfer N to host liverworts from organic sources (Field *et al*. unpublished), all point to an important role of Mucoromycotina FRE in host plant N nutrition. Indeed, our cytological analyses show that, differently from *Lycopodiella* roots where only fine endophytes were observed (Figure 2; Table 1), all other co-occurring plants (*F. foveolata, J. bulbosus, M. caerulea*) were also colonised by coarse endophytes with cytology typical of Glomeromycotina (Table 1). The finer functional details, in terms of N and P transfer, of this partnership in other vascular plants from a broader range of habitats remain to be established; the challenge here will be to separate the nutritional contributions of Mucoromycotina FRE and Glomeromycotina to host plants that are co-colonized by both fungi, as it seems to be the prevailing condition in vascular plants, especially angiosperms.

**Table 1.** Cytology of colonisation and fungal identity of study species (*) compared to relevant examples from the literature. (referred to in Discussion). *Results from this study; G = gametophyte generation; S = sporophyte generation; ICSs = intercellular spaces

|  | G/<br>S | Tissue/location | Colonization | Morphology (diameter) | Fungus<br>ID | References |
| --- | --- | --- | --- | --- | --- | --- |
| **Liverworts** |  |  |  |  |  |  |
| *Treubia*<sup>1</sup> | G | several ventral cell layers | intracellular | coils (0.5-1.5 µm) with 'lumps'/swellings (up to 15 µm), arbuscule-like short-side branches on coiled hyphae | M<sup>2</sup> | <sup>43</sup> Duckett *et al.* 2006<br><sup>5</sup> Bidartondo *et al.* 2011 |
|  |  | above intracellular zone | intercellular: large mucilage-filled ICSs | coarse hyphae 2-3 µm, thick-walled fungal structures |  |  |
| *Fossombronia* * | G | thallus central strand | intracellular | coarse hyphae (2-3 µm); large vesicles (15-30 µm), coils (0.5-1 µm), fine hyphae (0.5-1.5 µm) with small swellings/vesicles (5-10 µm), arbuscules | (M&G)* |  |
| **Lycophytes** |  |  |  |  |  |  |
| *Lycopodiella* * | G | outer cortical cell layers | intracellular | coils (up to 2.5µm) with vesicles (15-20µm), fine hyphae (0.5-1.5 µm) with small swellings/vesicles (5-10 µm) | M* |  |
|  |  | several ventral cell layers | intracellular |  |  |  |
|  |  | above intracellular zone | intercellular: large mucilage-filled ICSs | coarse hyphae (2- >3 µm) |  |  |
|  | S | protocorm: several ventral cell layers<br>central, above intracellular zone | intracellular<br>intercellular: large mucilage-filled ICSs | fine hyphae (0.5-1.5 µm) with small swellings/vesicles (5-10 µm)<br>coarse hyphae (2- >3 µm) | M* |  |
|  | S | root | intracellular and intercellular, small ICSs | fine hyphae (0.5-1.5 µm) with small swellings/vesicles (5-15 µm) | M<sup>1</sup>* | <sup>9</sup> Rimington *et al.* 2015 |
| **Angiosperms** |  |  |  |  |  |  |
| *Holcus* * | S | root | intracellular and intercellular, small ICSs | coarse hyphae (>3 µm), large vesicles (20-40 µm), fine hyphae (0.5-1.5 µm) with small vesicles (5-10 µm), arbuscules/arbuscule-like structures | (M&G)* |  |
| *Molinia* * | S | root | intracellular and intercellular, small ICSs | coarse hyphae (>3 µm), large vesicles (20-40 µm), fine hyphae (0.5-1.5 µm) with small vesicles/swellings (5-10 µm), arbuscules/arbuscule-like structures | (M&G)* |  |
| *Juncus* * | S | root | intracellular and intercellular, small ICSs | coarse hyphae (>3 µm), large vesicles (20-40 µm), fine hyphae (0.5-1.5 µm) with small vesicles (5-10 µm), arbuscules/arbuscule-like structures | (M&G)* |  |
| *Trifolium*<sup>1</sup> | S | root | intracellular and intercellular, small ICSs | coarse hyphae (>3 µm), large vesicles (>30 µm) fine hyphae (>1.5 µm), intercalary and terminal vesicles/swellings (5-10 µm) and arbuscules/arbuscule-like structures | (M&G)<sup>1</sup> | <sup>15</sup> Orchard *et al.* 2017 |
| **Fossils** |  |  |  |  |  |  |
| *Horneophyton*<sup>1</sup> | S | aerial axes, cortical cells | intracellular | coarse hyphae (>3 µm), large vesicles (up to 50 µm), arbuscule-like structures | G<sup>1</sup> | <sup>6</sup> Strullu-Derrien *et al.* 2014 |
|  |  | corm | intracellular and intercellular | intracellular coils, intercellular coarse hyphae (11-13 µm), thick-walled fungal structures | M<sup>1</sup> |  |
| *Nothia*<sup>1</sup> | S | aerial and prostrate axes | intercellular and intracellular | coarse hyphae (up to 15 µm) and intercellular vesicles (>50 µm) | ? | <sup>12</sup> Krings *et al.* 2007 |

### Mucoromycotina fine root endophytes

Mucoromycotina fungi within Endogonales colonising the gametophytes of liverworts (*F. foveolata*) and lycophytes (*L. inundata*), the sporophytic protocorms and roots of lycophytes (*L. inundata*) and the roots of angiosperms (*J. bulbosus, M. caerulea, H. lanatus*), all display the same characteristic morphology attributed previously to FRE [16, 19]. This contrasts with that typical of Glomeromycotina fungal associations, consisting of coarse hyphae (>3 µm diameter) and larger vesicles, which we observed in *Fossombronia, Juncus, Molinia, Holcus* but not in *L. inundata (*Table 1). These observations together with molecular identification of Mucoromycotina clades shared by these phylogenetically distant plant lineages support previous suggestions that vascular plants’ FRE are closely related to the Mucoromycotina mycorrhizal-like symbionts of non-vascular plants [15]. Here, we show that the same Mucoromycotina FRE are symbiotic across different land plant phyla.

Our demonstration of an extensive intercellular phase of fungal colonisation in the gametophytes and protocorms of *L. inundata* is in line with other lycophytes [10, 36] and strongly recalls the gametophytes of the Haplomitriopsida liverwort *Treubia* [31] and several hornworts [9], all of which have also been shown to associate with Mucoromycotina fungi [6, 9]. Differently from their fine intracellular counterparts, intercellular hyphae become swollen, eventually reaching more than 3 µm in diameter. Tightly wound hyphal coils up to 2.5 µm in diameter with somewhat larger terminal vesicles (up to 20 µm in diameter) are also prominent in the outer cortical layers of *L. inundata* gametophytes but were not observed in either protocorms or roots. Thus, Mucoromycotina FRE display considerable phenotypic plasticity in their interactions with ancient lineages of land plants which appears to relate to the developmental stage of the host and whether it produces an extensive network of mucilage-filled intercellular spaces. Comparable intercellular proliferation patterns alongside intracellular fungal structures have been described in the Devonian fossil plant *Horneophyton ligneri* and attributed to Mucoromycotina [7], closely resembling the distinctive inter- and extracellular fungal colonisation of another Devonian fossil plant, *Nothia* [47] (Table 1). The putative occurrence of Mucoromycotina FRE in early land plants and their presence in both extant early and later diverging plant lineages now point to a prominent role of these fungi, not only in plant terrestrialization [14], but also in current ecosystem functioning. Indeed, Mucoromycotina FRE have been shown to occur worldwide across many ecosystems, particularly in the roots of crop and pasture species, where colonization levels may be high, even as dense as the biomass of coarse Glomeromycotina arbuscular mycorrhizal fungi [19].

### More ammunition for the mycorrhizal revolution

Our findings provide, for the first time, conclusive evidence that Mucoromycotina FRE form nutritional mutualisms not only with non-vascular liverworts [5, 18] but also with a vascular plant. In line with previous reports showing nutritional mutualisms between non-vascular plants and Mucoromycotina fungi, with the exception of *Treubia lacunosa* [5, 8, 18], our experimental system was not significantly affected by a[CO_2_] concentrations. However, we report that *L. inundata* transfers up to 189 times more photosynthate to Mucoromycotina fungi than a non-vascular plant [5, 18]. In addition, we found that Mucoromycotina fungi transfer less ^33^P tracer, but can transfer ca. 391 times more ^15^N tracer to a vascular than to a non-vascular plant [18]. In contrast, the literature on FRE so far has focused on P [19]. From an evolutionary standpoint, our findings point towards a new physiological niche for the persistence of Mucoromycotina fungi from ancient to modern plants, both singly and in dual colonisation with Glomeromycotina.

## Acknowledgements

We gratefully acknowledge support from the NERC to KJF, SP, SS (NE/N00941X/1) and MIB (NE/N009665/1). KJF is funded by BBSRC Translational Fellowship (BB/M026825/1). WRR is supported by a NERC DTP (Science and Solutions for a Changing Planet) studentship. We thank James Giles (Natural England) for field support, Julia Masson and the RSPB for access to the Norfolk site, and The Species Recovery Trust for access to the Dorset site.

## Author’s contributions

K.J.F., S.P., S.S., M.I.B., and J.G.D. conceived and designed the investigation. S.P., J.K., J.G.D., M.I.B., A.S.J. and G.A.H collected plant material. G.A.H. and K.J.F. undertook the physiological analyses. A.S.J., W.R.R. and M.I.B. undertook the molecular anlayses. S.P. undertook the cytological analyses with assistance from J.K. P.G. and G.G. analysed and interpreted the stable istope data. All authors discussed results and commented on the manuscript.

