## Supplemental information for "Mucoromycotina fine root endophyte fungi form nutritional mutualisms with vascular plants"

**This PDF file includes:**

Supplementary text

Figs. S1 to S11

Tables S1 to S3

References for SI reference citations

**Other supplementary materials for this manuscript include the following:**

Supplementary Information Text

**Summary**

This file provides specific details on methods referred to in the main text of the above article, including mesh-covered core construction, equations for calculation of carbon and nutrient fluxes between symbionts and molecular methods for fungal identification. Cytological analyses of resin-embedded plant material are also included. Figures S1 and S9 illustrate experimental systems; Figures S2 to S8 are phylogenetic trees of Endogonales (Mucoromycotina). Figures S10 and S11 provide additional light and scanning electron micrographs of targeted plant species colonized by Mucoromycotina fine root endophytes (FRE), which complement those in the main text.

**Materials and Methods**

**Plant growth conditions**

*Lycopodiella inundata* plants were maintained in controlled environment chambers (Micro Clima 1200, Snijders Labs, Netherlands) with settings chosen to simulate the plant’s natural environment: 70% RH, 16 hr/8 hr day/night, 20°C day/ 15°C night, 100.1 µmol m^-2^ s^-1^ irradiance.

**Cytological analysis**

For trypan blue staining, plant tissues were cleared in 5% KOH at 90^o^C for 3 h, acidified in 2% HCl for 1-2 min, stained in 0.05% trypan blue, de-stained in 50% glycerin for 24 h before viewing under a light microscope. For SEM, tissues were fixed in 3% glutaraldehyde, dehydrated through an ethanol series, critical-point dried using CO_2_ as transfusion fluid, sputter coated with 390 nm palladium-gold and viewed using a FEI Quanta scanning electron microscope (FEI, Hillsboro, OR, USA).

For light microscopy, *L. inundata* gametophytes and the protocorms of young sporophytes were processed according to Ligrone and Duckett [4]. Briefly, specimens were fixed in 3% glutaraldehyde, 1% freshly prepared formaldehyde and 0.75% tannic acid in 0.05 M Na–cacodylate buffer, pH 7, for 3 h at room temperature. After several rinses in 0.1 M buffer, samples were post-fixed in buffered (0.1 M, pH 6.8) 1% osmium tetroxide overnight at 48^o^ C, dehydrated in an ethanol series and embedded in Spurr’s resin via ethanol, and 0.5 mm thick sections were cut with a diamond histo-knife, stained with 0.5% toluidine blue and photographed under a Zeiss Axioscope light microscope equipped with an MRc digital camera

**^33^P sample analysis**

After harvest and freeze-drying, 10-50 mg of homogenised plant and soil materials were digested in 500 µl of concentrated H_2_SO_4_. These were heated to 365°C for 15 min, and 50 µl H_2_O_2_ were added to each sample when cool. Samples were reheated to 365°C for one minute, producing a clear digest solution which was then cooled and diluted to 5 ml with distilled water. One ml of each diluted digest was then added to 10 ml of the scintillation cocktail Emulsify-safe (Perkin Elmer, Beaconsfield, UK) before liquid scintillation counting.

**^14^C sample analysis**

Approximately 10-20 mg of scintillation of sample was weighed into Combusto-cones (Perkin Elmer) which were then oxidised. CO_2_ released through oxidation was trapped in 10 ml Carbosorb (Perkin Elmer, UK) prior to mixing with the scintillation cocktail 10 ml Permaflour (Perkin Elmer, UK). Total carbon (^12^C + ^14^C) fixed by the plant and transferred to the fungal network was calculated as a function of the total volume and CO_2_ content of the labelling chamber and the proportion of the supplied ^14^CO_2_ label fixed by plants (see SI). The difference in carbon between the static and rotated cores is considered equivalent to the total C transferred from plant to symbiotic fungus within the soil core, noting that a small proportion will be lost through soil microbial respiration. The total carbon budget for each experimental pot was calculated using equations from Cameron *et al*. [3].

**Stable isotope signatures of neighbouring plants**

Leaf and root samples were washed, dried to constant weight at 105°C and ground to a fine powder in a ball mill (Retsch Schwingmühle MM2, Haan, Germany). Relative C and N isotope natural abundances of the leaf and root samples were measured in a dual element analysis mode with an elemental analyser (Carlo Erba Instruments 1108, Milan, Italy) coupled to a continuous flow isotope ratio mass spectrometer (delta S, Finnigan MAT, Bremen, Germany) via a ConFlo III open-split interface (Thermo Fisher Scientific, Bremen, Germany) as described in Bidartondo *et al*. [3]. Relative isotope abundances are denoted as δ values calculated according to the following equation: δ^13^C or δ^15^N = (*R*_sample_/*R*_standard_ – 1) x 1000 [‰], where *R*_sample_ and *R*_standard_ are the ratios of heavy to light isotope of the samples and the respective standard. Standard gases were calibrated with respect to international standards (CO_2_ vs PDB, N_2_ vs N_2_ in air) with the reference substances ANU sucrose and NBS18 for the carbon isotopes and N1 and N2 for the nitrogen isotopes (International Atomic Energy Agency, Vienna, Austria). Reproducibility and accuracy of the C and N isotope abundance measurements were routinely controlled by measuring the laboratory standard acetanilide. In relative C and N isotope natural abundance analyses, acetanilide was routinely analyzed with variable sample weight at least six times within a batch of 50 samples. The maximum variation of δ^13^C and δ^15^N was always below 0.2‰.

**Construction of mesh-covered cores**

Based on the methods of Field *et al*. [1] two windows (20 mm x 50 mm, Supplementary Figure S1) were cut into the below-ground portion of each core. The windows and base were covered with nylon mesh (10 µm pore size) and sealed with a fast-setting acrylic adhesive (Tensol 12, Bostok Limited, UK). This mesh size is fine enough to exclude lycophyte roots but allows the ingrowth of fungal hyphae. We perforated a fine-bore capillary tube using a needle, and installed it to run the full-length of each of the cores (100 mm in length, 1.02 mm internal diameter; Portex, UK). The tubing ensured isotope solution was introduced and distributed evenly throughout the core volume. We sealed the capillary tube using acrylic adhesive 5 mm from the bottom of the core in order to prevent excess isotope leaching from the bottom of the core.

**Sampling of lycophytes, liverworts and angiosperms**

Sampling sites for *Lycopodiella inundata* and adjacent liverworts and angiosperms are listed in Supplementary Table 1 (S1).

**Plant harvest and sample analyses**

Upon detection of maximal belowground ^14^C flux following release of ^14^CO_2_, 3ml of 2 M KOH was introduced to each chamber to trap residual ^14^CO_2_ gas in the chamber headspace. One ml of each ‘used’ KOH trap was transferred to vials containing 10 ml of the scintillation cocktail Ultima Gold (Perkin Elmer, Beaconsfield, UK) and the radioactivity of each sample determined through liquid scintillation. These data were used to calculate carbon budgets for each experimental pot (see below for equations).

Plant and soil materials were separated, freeze-dried, weighed and homogenised using a Tissue Lyser LT (Qiagen, UK).

**^33^P transfer from fungus to plant**

The ^33^P transferred from fungus to plant was determined using equation 1 [2]:

(1) *M*^33^P = {$[\frac{A}{S\mathrm{Act}}]Mwt\} Df$

Where *M*^33^P = Mass of ^33^P, *A =* radioactivity of the tissue sample (Bq), *S*Act = specific activity of the source (Bq mmol^-1^), *Df* = dilution factor and *M*wt = molecular mass of P.

**Carbon transfer from plant to fungus**

The difference in carbon between the static and rotated cores is taken to be equivalent to the total C transferred from plant to symbiotic fungus within the soil core. Total carbon assimilated by the plant was calculated using equation 2 [4]:

(2) *M_C_* = $\left( \left( \frac{A}{S\mathrm{Act}} \right)M14C \right)+(P_{r}x {M\mathrm{wt}}_{c})$

Where *M_C_* = Mass of carbon transferred from plant to fungus, *A* = radioactivity of the tissue sample (Bq); *S*Act = specific activity of the source (Bq Mol^-1^), *M^14^C* = atomic mass of ^14^C, *P*_r_ = proportion of the total ^14^C label supplied present in the tissue; *M*wt_C_ = mass of C in the CO_2_ present in the labelling chamber (g) (from the ideal gas law; Equation 3):

(3) *M_cd_* = *M_cd_* $\binom{{PV}_{CD}}{RT}\therefore m_{c = m_{cd}}$x 0.27292

Where *m_cd ­_=* mass of CO_2_ (g), *M_cd_ =* molecular mass of CO_2_ (44.01 g mol^-1^) P = total pressure (kPa); *V_cd_* = volume of CO_2_ in the chamber (0.003 m^3^); *R* = universal gas constant (J K^-1^ mol^-1^); *T*, absolute temperature (K); *m_c_,* mass of C in the CO_2_ present in the labelling chamber (g), where 0.27292 is the proportion of C in CO_2_ on a mass fraction basis [3]

**Suppliers and addresses**

Petersfield no.2 compost, Leicester, UK

Snijder Labs, Netherlands

Vaisala, Birmingham, UK

Zeiss, Germany

Sigma, UK

Hartmann Analytics, Germany

Isotech, Chesterfield, UK

Sercon Ltd., Crewe, UK

Carlo Erba Instruments, Milan, Italy

delta S, Finnigan MAT, Bremen, Germany

Thermo Fisher Scientific, Bremen, Germany

IBM Analytics, New York, USA

**Perforated capillary tube**

**90 mm**

**20 mm**

**10 µm mesh**

Figure. S1. Schematic diagram of mesh-covered core (not drawn to scale).

**Table S1. Samples of lycophytes, liverworts and angiosperms analysed with their origin.**

| **Sample species** | **Samples analysed** | **Origin** |
| --- | --- | --- |
| **Lycophytes** |  | |
| *Lycopodiella inundata* (L.) Holub | 24 | Thursley Common, Surrey; England |
| *Lycopodiella inundata* (L.) Holub | 16 | Studland Heath, Dorset; England |
| *Lycopodiella inundata* (L.) Holub | 4 | confidential site, Norfolk; England |
| **Liverworts** |  | |
| *Fossombronia foveolata* Lindb. | 1 | Thursley Common, Surrey; England |
| *Fossombronia foveolata* Lindb. | 3 | confidential location, Norfolk; England |
| *Fossombronia foveolata* Lindb. | 6 | Lynn Crafnant; Wales |
| **Angiosperms** |  | |
| *Molinia caerulea* (L.) Moench | 28 | Thursley Common, Surrey; England |
| *Juncus bulbosus* L. | 30 | Thursley Common, Surrey; England |
| *Juncus bulbosus* L. | 3 | confidential location, Norfolk; England |
| *Holcus lanatus* L. | 3 | Lynn Crafnant; Wales |

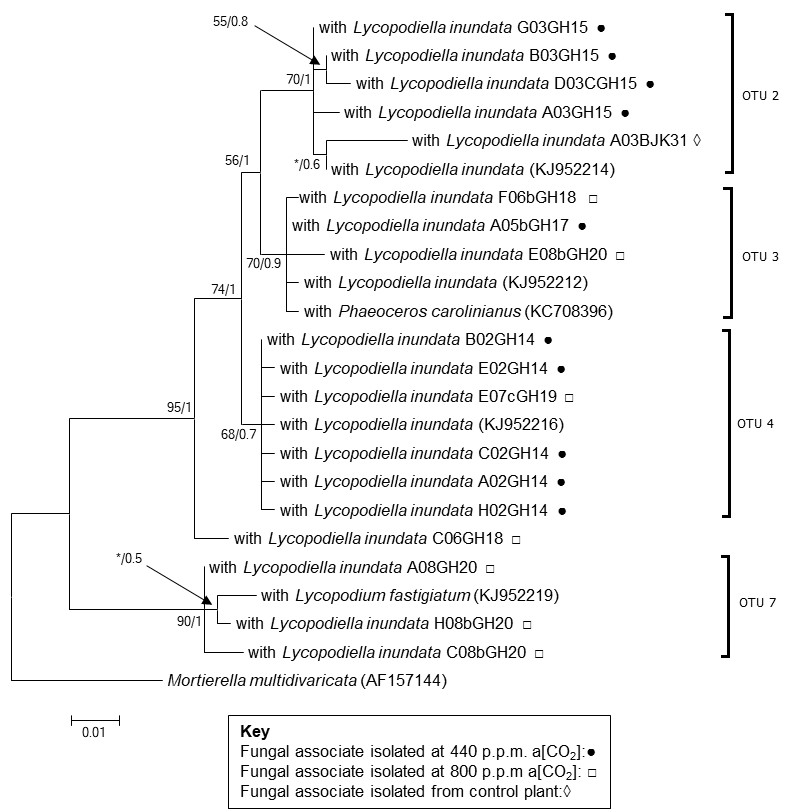

**Figure S2. Phylogenetic relationships of Mucoromycotina OTUs associated with *Lycopodiella inundata* grown at ambient (440 ppm a[CO_2_]) and elevated (800 ppm a[CO_2_]) conditions. The final alignment consisted of 24 taxa and 397 characters. The TN93 + G model was used for analysis. The ML tree is shown. The values adjacent to each node correspond to the bootstrap support and posterior probabilities, respectively. An asterisk denotes a bootstrap value <50% or posterior probability <0.5.**

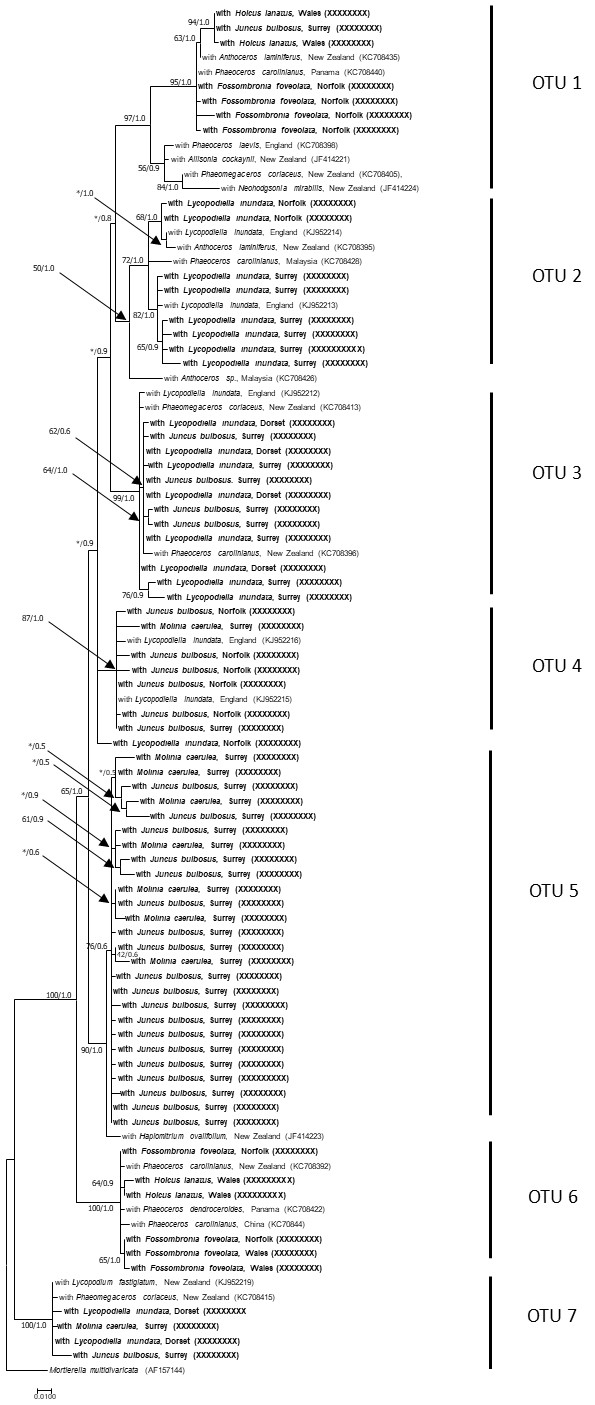

**Figure S3. An overview of phylogenetic relationships of Mucoromycotina OTUs associated with bryophytes, lycophytes and angiosperms from various UK locations based on partial 18S gene sequences. Only the subset of representative OTUs relevant to this study are shown. Our sequences lie within the *Densosporaceae* which is sister to the *Endogonaceae* (Desirò *et al.* [6]). The final alignment consisted of 94 taxa and 693 characters. Sequences shorter than 690 bp were excluded from the analysis. The model selected for analysis was K80 + G + I. The tree shown is that inferred from the ML analysis. The values adjacent to each node correspond to the bootstrap support and posterior probabilities, respectively. An asterisk denotes a bootstrap value <50% or posterior probability <0.5.**

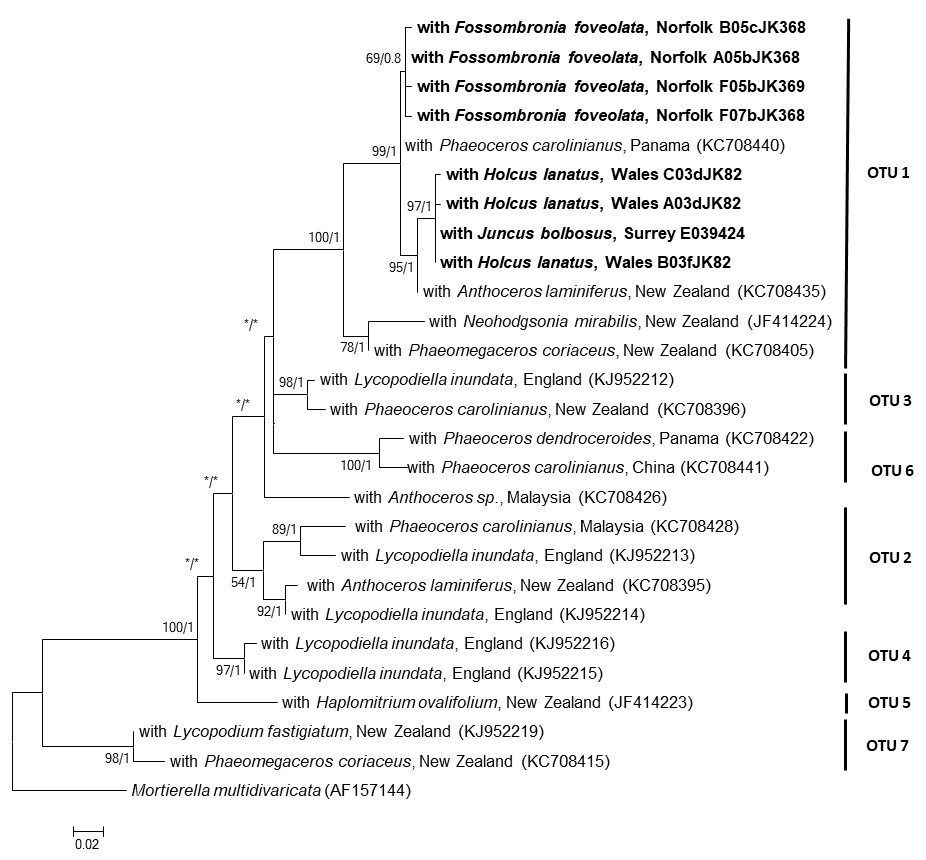

**Figure S4. Phylogenetic relationships of partial 18S DNA sequences classified as OTU 1, corresponding to “group A” in Desirò *et al.* [6]. The ML tree is shown. Values at nodes refer to ML/Bayesian inference. The final alignment consisted of 731 characters and analysis was conducted using the TN93 + G + I substitution model.**

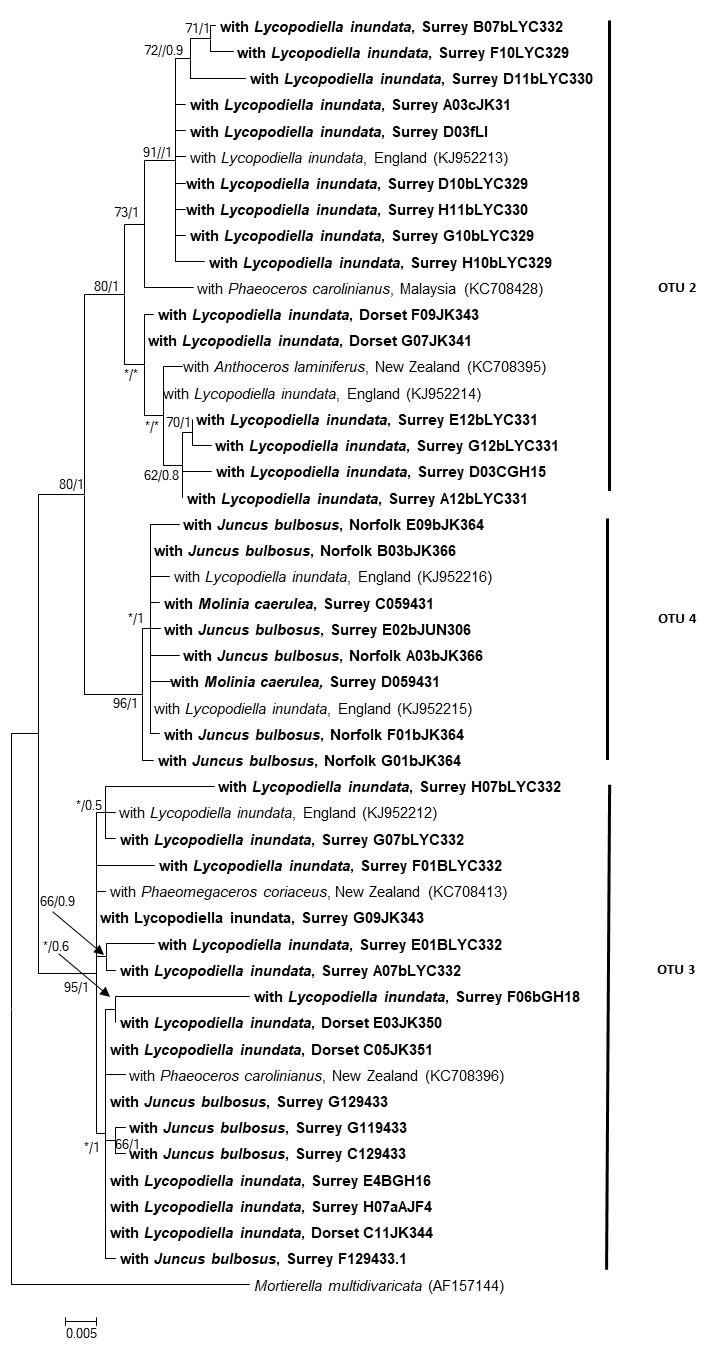

**Figure S5.** **Phylogenetic relationships of partial 18S DNA sequences classified as OTU 2-4. The ML tree is shown. The final alignment consists of 681 characters and phylogenetic analysis was conducted using the best fit model, TN93 + G, as explained in materials and methods except that 2,000,000 generations were run for Bayesian analysis. The values adjacent to each node correspond to the bootstrap support and posterior probabilities, respectively. An asterisk denotes a bootstrap value <50% or posterior probability <0.5.**

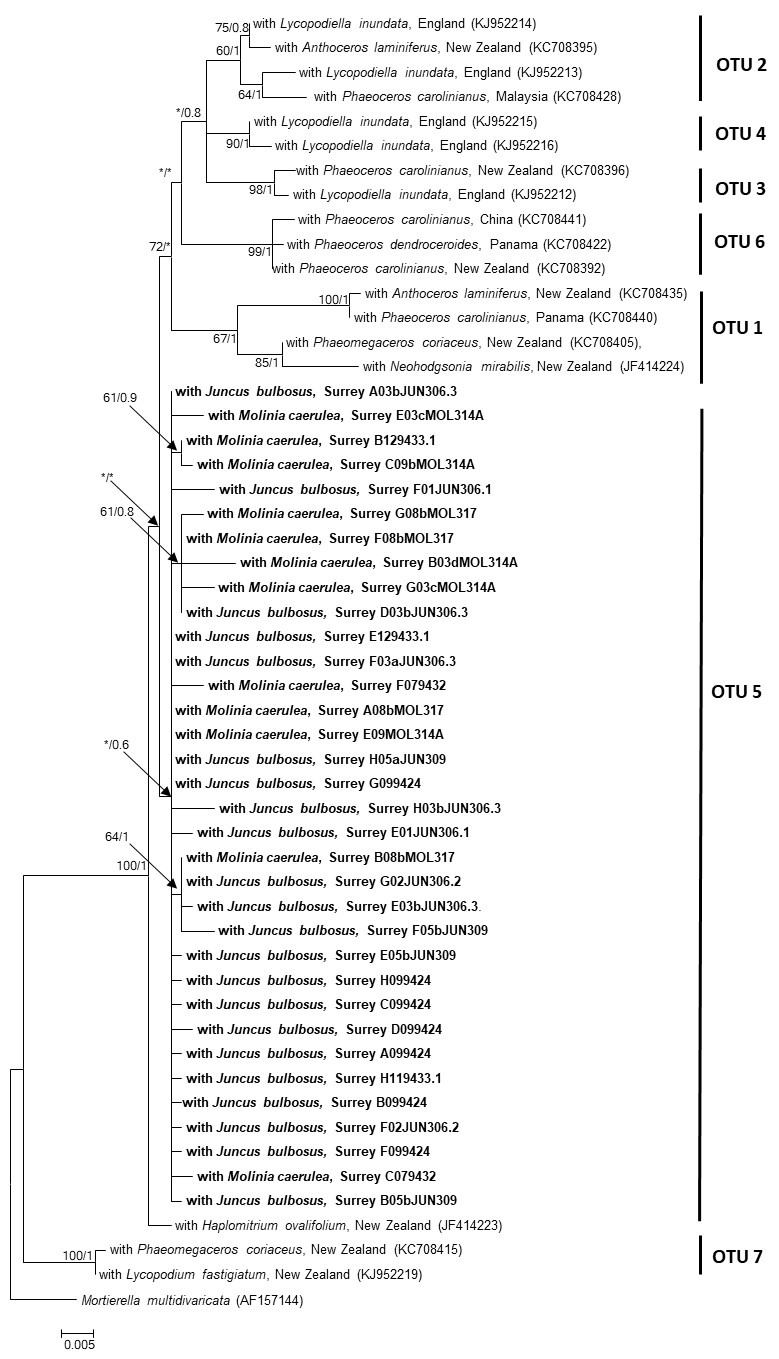

**Figure S6**. **Phylogenetic relationships of partial 18S DNA sequences clustering within OTU 5. The final alignment comprised 649 characters and used the TN93 +G substitution model. The ML tree is featured. The values adjacent to each node correspond to the bootstrap support and posterior probabilities, respectively. An asterisk denotes a bootstrap value <50% or posterior probability <0.5.**

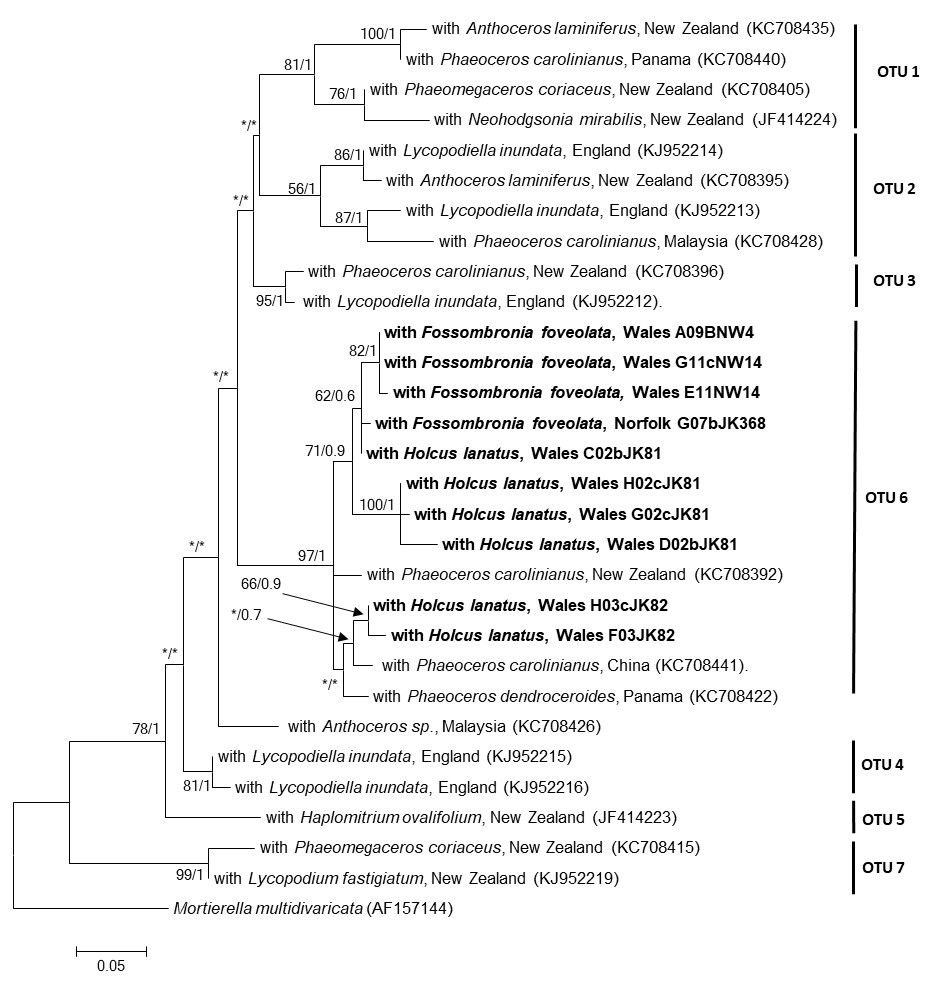

**Figure S7**. **Phylogenetic relationships of partial 18S DNA sequences classified as OTU 6, corresponding to “group B” in Desirò *et al*. [6]. The ML tree is displayed. The final dataset consisted of 622 characters and was analysed using the TN93 + G + I substitution model. Values at the nodes indicate ML/Bayesian inference values.** **An asterisk denotes a bootstrap value <50% or posterior probability <0.5.**

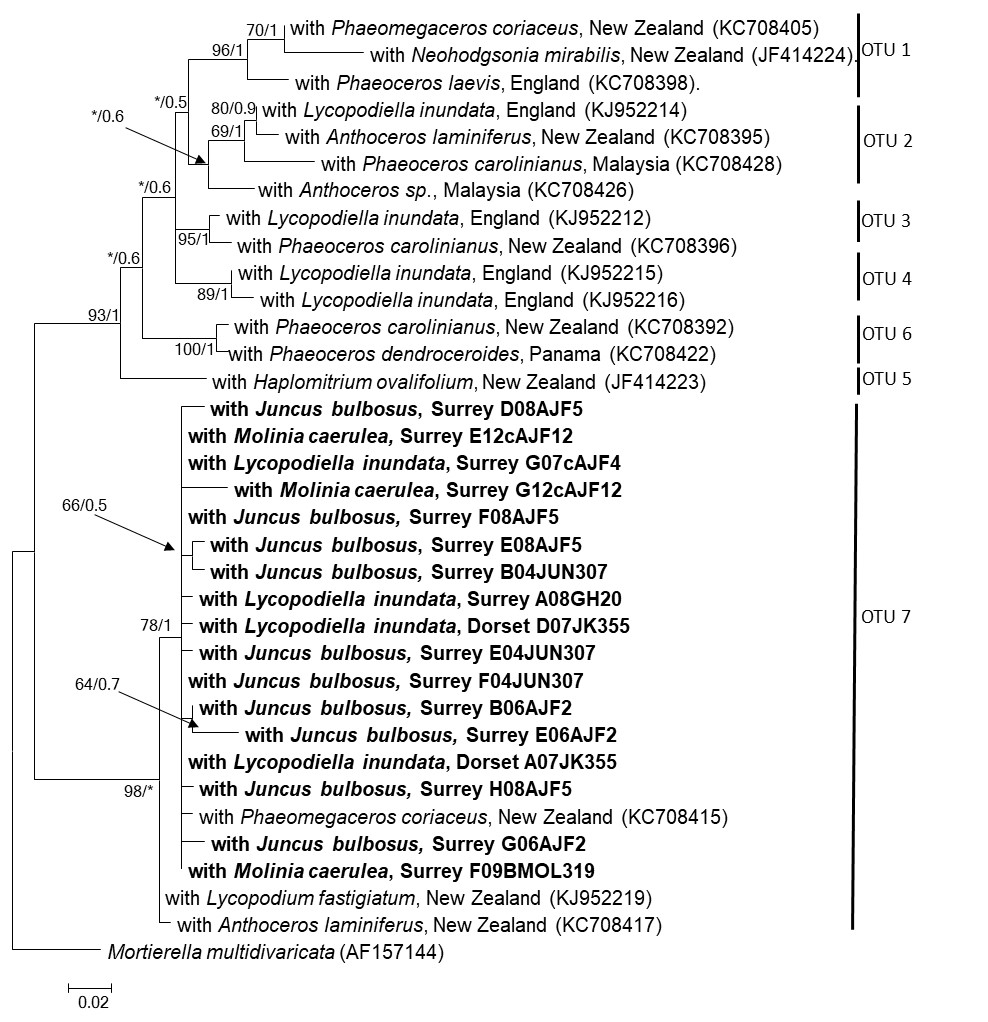

**Figure S8**. **Phylogenetic relationships of partial 18S DNA sequences clustering with OTU 7, corresponding to “group I” in Desirò *et al.* [6]. The ML tree is shown. The final alignment comprised 580 characters and the substitution model used for the analysis was HKY85 + G + I. Values at the nodes indicate ML/Bayesian inference values. An asterisk denotes a bootstrap value <50% or posterior probability <0.5.**

**A**

**^15^N**

**^33^P**

**^15^N**

**^33^P**

**B**

**^14^CO_2_**

**^14^CO_2_ fixed through photosynthesis**

**^14^C sugars and/or lipids transferred to fungus within core**

**^14^C Sodium bicarbonate**

**10% Lactic Acid**

Figure S9. Illustration of experimental procedure for labelling lycophytes with (A) ^33^P-orthophosphate and ^15^N and (B) ^14^CO_2_.

**Table S2**. A summary of Mucoromycotina OTUs associated with liverworts, lycophytes and angiosperms at four UK sites. The numbers within each column represent the number of partial 18S sequences that cluster within each OTU.

| **Species** | **Location** | **OTU 1** | **OTU 2** | **OTU 3** | **OTU 4** | **OTU 5** | **OTU 6** | **OTU 7** | **ND^1^** |
| --- | --- | --- | --- | --- | --- | --- | --- | --- | --- |
| *Lycopodiella inundata*  (*n* = 4) | Dorset | 0 | 2 | 4 | 0 | 0 | 0 | 3 | 0 |
| *Fossombronia foveolata*  (*n* = 2) | Norfolk | 4 | 0 | 0 | 0 | 0 | 1 | 0 | 0 |
| *Juncus bulbosus*  (*n* = 2) | Norfolk | 0 | 0 | 0 | 5 | 0 | 0 | 0 | 0 |
| *Lycopodiella inundata*  (*n* = 1) | Norfolk | 0 | 0 | 0 | 0 | 0 | 0 | 0 | 1 |
| *Lycopodiella inundata*  (*n* = 6) | Surrey | 0 | 13 | 7 | 0 | 0 | 0 | 4 | 0 |
| *Juncus bulbosus*  (*n* = 12) | Surrey | 1 | 0 | 4 | 13 | 31 | 0 | 16 | 0 |
| *Molinia caerulea*  (*n* = 8) | Surrey | 0 | 0 | 0 | 9 | 19 | 0 | 12 | 0 |
| *Fossombronia foveolata*  (*n* = 2) | Wales | 0 | 0 | 0 | 0 | 0 | 4 | 0 | 0 |
| *Holcus lanatus*  (*n* = 2) | Wales | 6 | 0 | 0 | 0 | 0 | 8 | 0 | 0 |

**^1^ND = not determined, singleton OTU.**

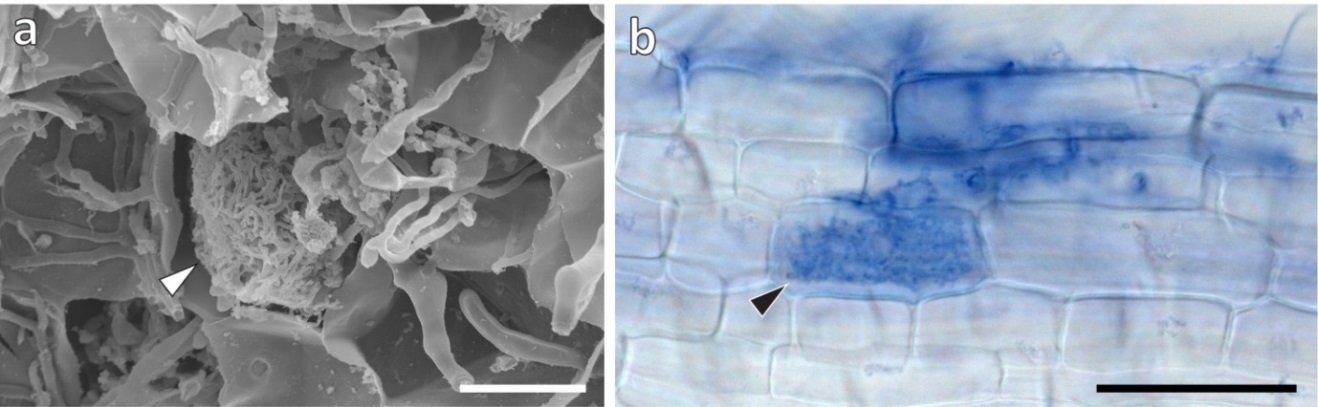

**Figure. S10** (a) Scanning electron micrograph of *Fossombronia foveolata* thallus showing coil of fine hyphae (arrowhead) and coarse hyphae in surrounding cells; (b) Light micrograph of trypan blue stained root of *Holcus lanatus* showing fine hyphae and an arbuscule-like structure (arrowhead). Scale bars: (b) 50 µm, (a) 20 µm.

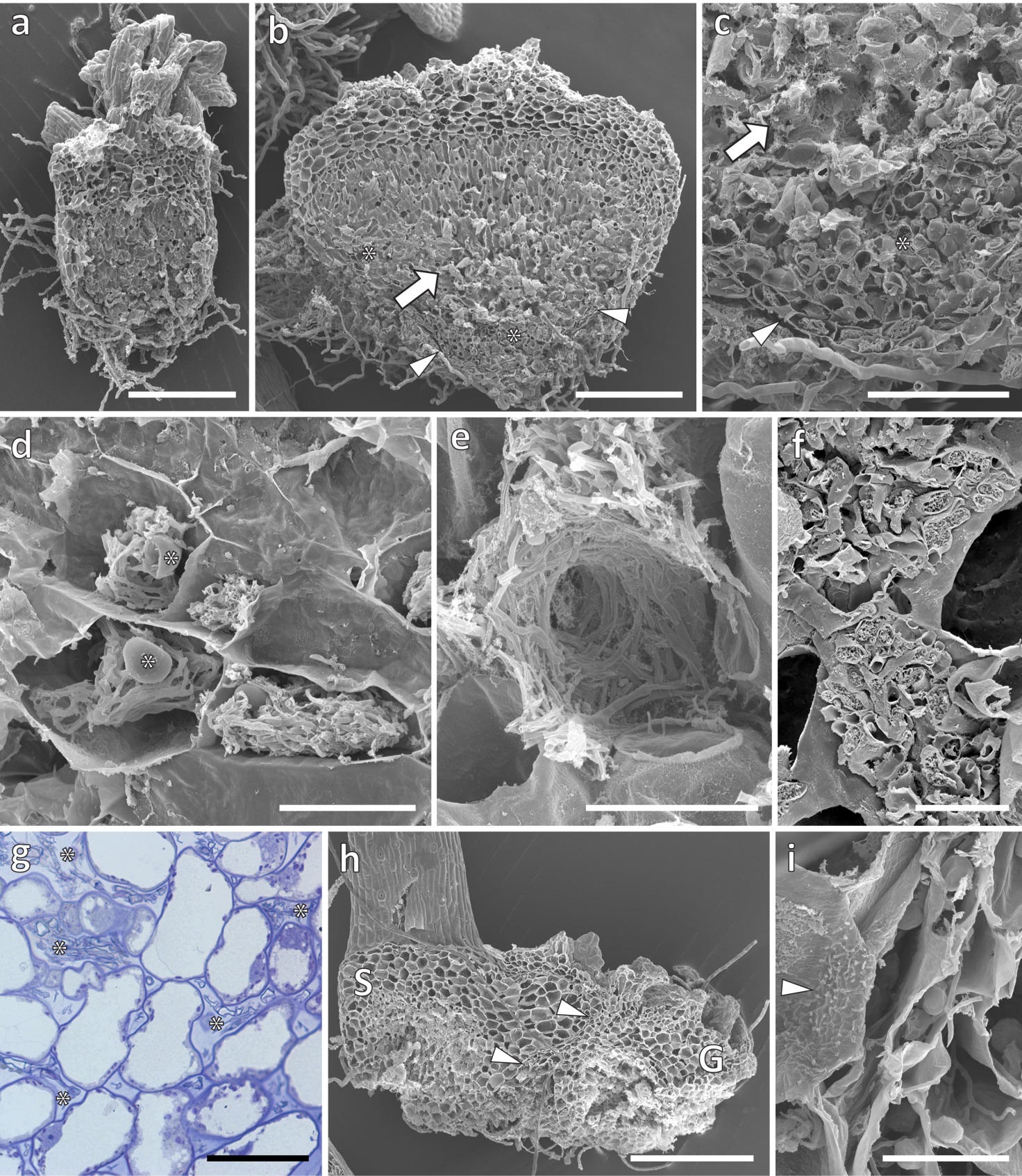

**Figure S11:** **Scanning electron micrographs, except (g) light micrograph of toluidine blue stained semi-thin sections. Gametophyte morphologies in *Lycopodiella inundata* (a, b), (b, c) zonation of fungal colonization in a gametophyte: intracellular in the outer cortical layers (arrowheads) and surrounding tissue (*), and strictly intercellular in the central region characterized by large, mucilage-filled, intercellular spaces (arrows); (d) hyphal coils and vesicles (*) in the outer cortical layers; (e) centrally, the fungus colonizes the system of large intercellular spaces; here the fine hyphae (e) eventually become swollen, reaching diameters of > 3 µm (f); (g) intercellular fungal proliferation (*), note the non-colonized, living host cells in this zone; (h) young sporophyte (S) attached to gametophyte (G), arrowheads point to the sporophyte-gametophyte junction, enlarged in (i). Note that the gametophyte fungus does not cross the placenta identifiable by its numerous wall ingrowths (arrowhead). Scale bars: (a, b, h) 500 µm, (c) 200 µm, (d, g) 50 µm, (e, f, i) 20 µm.**

**Table S3. Summary of differences in Mucoromycotina functionality between non-vascular and vascular plants. Mucoromycotina functionality in *Haplomitrium gibbsiae* and *Treubia lacunosa* (non-vascular) and *Lycopodiella inundata* (vascular) at simulated Paleozoic a[CO_2_] conditions on Earth when these plants are thought to have diverged [8] and modern day ambient a[CO_2_] conditions. Values denote absolute values for C (ng), and concentrations of P & N (ng/g) in plant tissue across all experiments.**

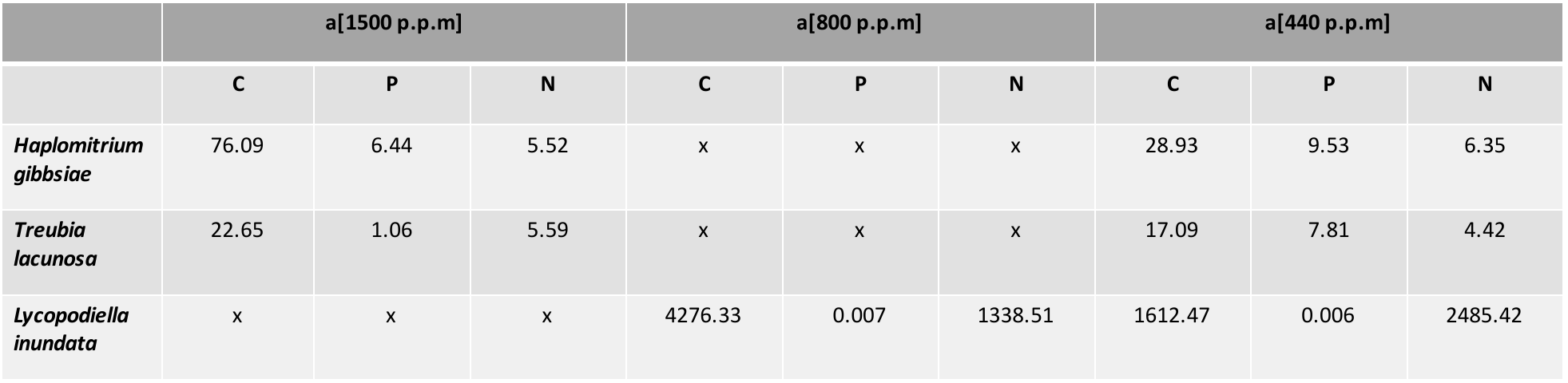
